# Responses to Temperatures of Different *Drosophila* Species

**DOI:** 10.1101/2021.10.01.462748

**Authors:** Ainul Huda, Thomas J. Vaden, Alisa A. Omelchenko, Allison N. Castaneda, Lina Ni

## Abstract

Temperature is a critical environmental variable that affects the distribution, survival, and reproduction of most animals. Although temperature receptors have been identified in different animals, how these receptors respond to temperatures is largely unknown. Here we use modified single-fly thermotactic assays to analyze movements and temperature preferences of nine *Drosophila* species. The ability/inclination to move varies among these species and at different temperatures. Importantly, different species prefer various ranges of temperatures. While wild-type *D. melanogaster* flies avoid the warm temperature in the warm avoidance assay and the cool temperature in the cool avoidance assay, *D. bipectinata* and *D. yakuba* avoid neither warm nor cool temperatures and *D. biarmipes* and *D. mojavensis* do not avoid the warm temperature in the warm avoidance assay. These results demonstrate that *Drosophila* species have different mobilities and temperature preferences, thereby benefiting the research on molecular mechanisms of temperature responsiveness.

**Summary statement:** The ability to move and the preference for temperatures vary among fly species when flies are exposed to steep temperature gradients.

## Introduction

Temperature affects all aspects of physiology, from the rate of chemical reactions and the activity of biomolecules to the distribution of living organisms (Dell et al., 2011, Sengupta and Garrity, 2013, Franks and Hoffmann, 2012). Temperature variation is particularly influential for small animals, such as insects, which depend on ambient temperatures to set their body temperatures (Garrity et al., 2010, Dillon et al., 2010). Many insect vectors of diseases, including mosquitoes, respond to the temperature of their warm-blooded hosts and use it to guide blood-feeding behaviors (Brown, 1951, Howlett, 1910, Corfas and Vosshall, 2015, Greppi et al., 2015, Greppi et al., 2020).

Fruit flies are a good insect model system to study thermosensation. In *D. melanogaster*, many thermosensory systems are governed by a small number of sensory neurons but control robust behaviors (Ni et al., 2013, Hamada et al., 2008, Klein et al., 2015, Budelli et al., 2019, Gallio et al., 2011). These sensory neurons possess evolutionarily conserved thermal molecules across *Drosophila* species and with insect vectors of diseases, including mosquitoes (Corfas and Vosshall, 2015, Greppi et al., 2020). Adult *D. melanogaster* flies possess several thermosensory systems to control different thermotactic behaviors (Barbagallo and Garrity, 2015). Aristal warm and cool neurons guide rapid warm and cool avoidance when flies are exposed to steep temperature gradients. A gustatory receptor GR28B(D) is the warm receptor in aristal warm neurons and three members from the Ionotropic Receptor (IR) family (IR25a, IR93a, and IR21a) form the cool receptor in aristal cool neurons (Ni et al., 2013, Budelli et al., 2019). However, molecular mechanisms underlying how these thermoreceptors respond to temperatures are largely unknown.

Besides *D. melanogaster*, genomes of more than 20 other *Drosophila* species have been sequenced (Celniker et al., 2002, Hoskins et al., 2007, Hu et al., 2013, Clark et al., 2007, Stark et al., 2007, Chen et al., 2014). These sequenced species span a wide range of global distributions with diverse temperatures (Powell, 1997). Therefore, they may possess thermoreceptors that have evolved distinct temperature responsiveness through amino-acid changes at a few residues to adapt to their specific ecosystems. We thereby expect that they will offer opportunities to understand how thermoreceptors respond to temperatures.

This study modifies single-fly thermotactic assays and uses TrackMate to track fly movement (Budelli et al., 2019, Tinevez et al., 2017). Using these assays, we test 801 flies from four genotypes of *D. melanogaster* and eight other *Drosophila* species. We find starting temperatures and genders affect preference indices and identify several *Drosophila* species mimicking temperature preferences of *D. melanogaster* thermoreceptor mutants. These results may benefit the research about molecular mechanisms of temperature responsiveness.

## Results

### *Drosophila* species have diverse mobilities

To understand temperature responses of different *Drosophila* species by thermotactic assays, we first analyzed their mobilities. Using single-fly warm and cool avoidance assays, we tested 801 flies, including four genotypes of *D. melanogaster* as controls, *D. ananassae, D. biarmipes, D. bipectinata, D. erecta, D. ficusphila, D. mojavensis, D. simulans*, and *D. yakuba*. A single fly was acclimated under a transparent cover (this cover is 83mm (length) X 58mm (width) X 2mm (height) and thus flies can only walk, but not fly) at 25±1°C for 2 min and then allowed to explore between zones of 25±1°C and 31±1°C or 11±1°C for 2 min. Their positions were recorded by TrackMate and moving distances were calculated. Next, we analyzed moving distances by pseudo-F statistics and identified 10 clusters. Flies in the cluster with the least moving distances moved from 107.584 to 799.243 pixels. Independent visual analysis by four researchers agreed that flies in this cluster had limited mobilities; flies in other clusters were able to adequately explore both temperatures (Movie 1). 800 pixels was set as the threshold (the dashed line in Fig. 1). Flies that had moving distances shorter than 800 pixels were omitted from preference index (PI) analysis.

**Fig 1.**
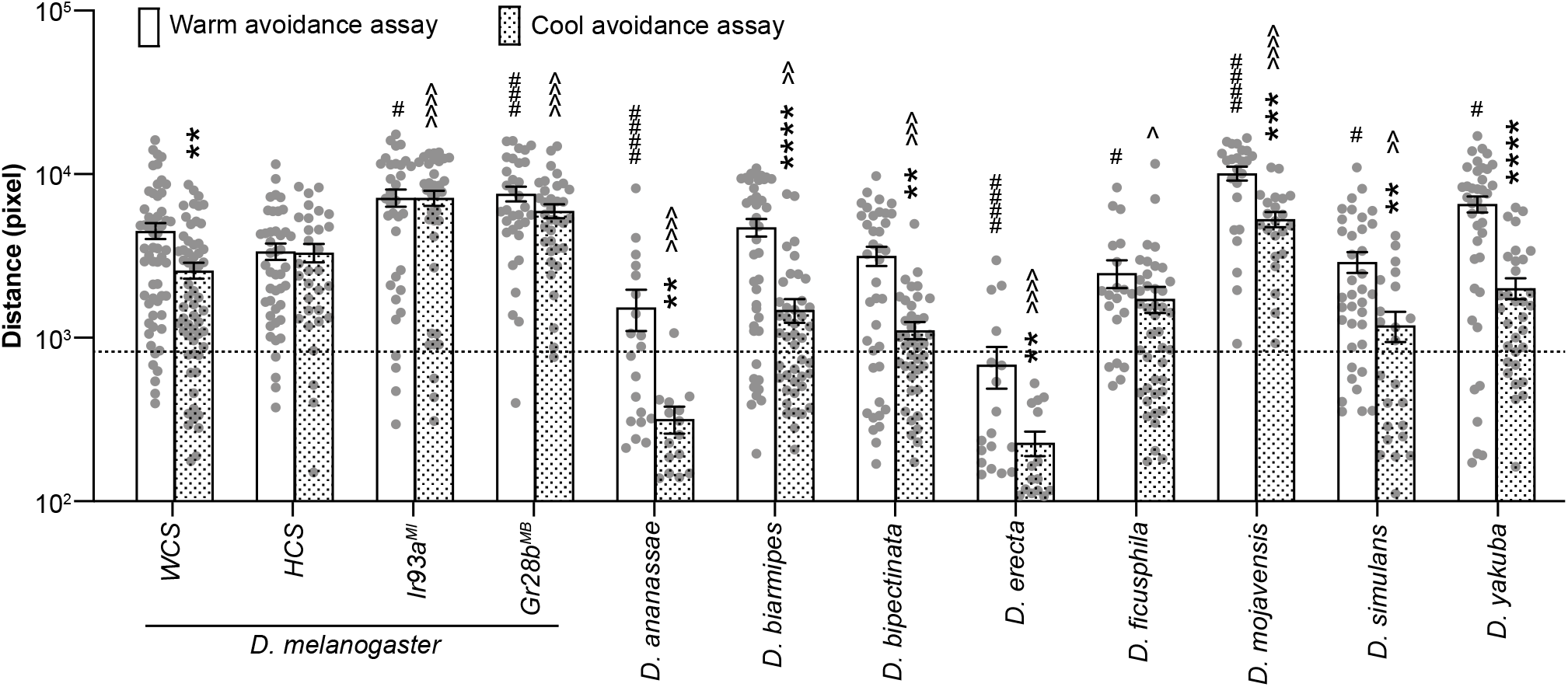
*Drosophila* species have diverse mobilities. Four genotypes (*WCS, HCS, Ir93a*^*MI*^, and *Gr28b*^*MB*^) of *D. melanogaster*, as well as *D. ananassae, D. biarmipes, D. bipectinata, D. erecta, D. ficusphila, D. mojavensis, D. simulans*, and *D. yakuba* were tested. The dashed line shows the threshold of the moving distance, 800 pixels. Data represent mean ± SEM. ** *p* < 0.01, *** *p* < 0.001, and **** *p* < 0.0001; comparing moving distances of the corresponding genotype/species in the warm avoidance assay; Mann-Whitney test, except Welch’s test for *Gr28b*^*MB*^. # *p* < 0.05, ### *p* < 0.001, and #### *p* < 0.0001; comparing moving distances of *WCS* in the warm avoidance assay; Mann-Whitney test. ^ *p* < 0.05, ^^ *p* < 0.01, ^^^ *p* < 0.001, and ^^^^ *p* < 0.0001; comparing moving distances of *WCS* in the cool avoidance assay; Mann-Whitney test.

Most fly species moved significantly more in the warm avoidance assay than in the cool avoidance assay, including *WCS D. melanogaster, D. ananassae, D. biarmipes, D. bipectinata, D. erecta, D. mojavensis, D. simulans*, and *D. yakuba* (Fig. 1). These data suggest that flies are more active in warm environments.

Moreover, fly species had diverse mobilities. In the warm avoidance assay, *D. erecta* moved the least and *D. mojavensis* moved the most. The average moving distance of *D. erecta* was about 1/15 of *D. mojavensis*. In the cool avoidance assay, *D. erecta* still moved the least, while the most active flies were *D. melanogaster Ir93a*^*MI*^, whose average moving distance was over 31 times that of *D. erecta* (Fig. 1). Of note, in the warm avoidance assay, moving distances from more than half of *D. ananassae* and *D. erecta* flies did not reach the threshold (Fig. 1) and were not further analyzed. Similarly, in the cool avoidance assay, less than half of *D. ananassae, D. erecta*, and *D. simulans* flies reached the threshold (Fig. 1) and their PIs were also not calculated.

### Starting temperatures and genders affect PIs

Next, we tried to understand the effects of starting temperatures and genders in PIs using wild-type *WCS D. melanogaster*. The PI was calculated by dividing the difference of the time spent at 25°C and the time spent at 31°C or 11°C by the total time. A positive PI indicates preference for 25°C, while a negative PI indicates preference for 31°C or 11°C; PI near zero suggests no preference.

We divided *WCS* data from the warm avoidance assay into four groups: males starting at 25°C, females starting at 25°C, males starting at 31°C, and females starting at 31°C. As shown in Fig. 2A, males and females had similar PIs when they started at the same temperatures. When flies started at different temperatures, PIs were significantly different. Flies that started at 25°C had strong preferences for 25°C, while flies that started at 31°C had no preferences between 25°C and 31°C. These data suggest that starting temperatures, but not genders, affect PIs in the warm avoidance assay.

**Fig 2.**
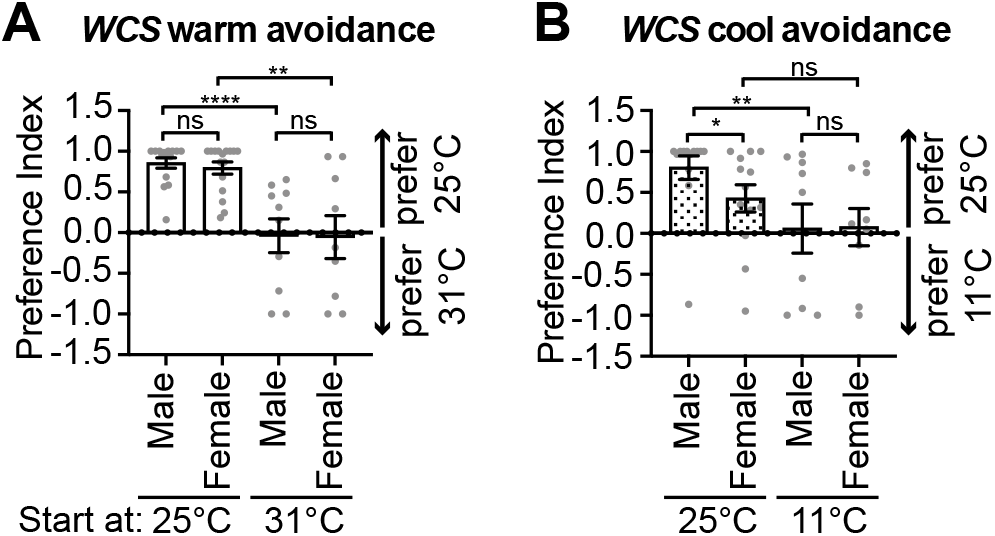
Both starting temperatures and genders affect *WCS* PIs. PIs of indicated groups in warm (A) and cool (B) avoidance assays. Data represent mean ± SEM. * *p* < 0.05, ** *p* < 0.01, and **** *p* < 0.0001; Mann-Whitney test, except Welch’s test for the comparison of males starting at 31°C and females starting at 31°C in (A) and the comparison of females starting at 25°C and females starting at 11°C in (B).

In the cool avoidance assay, we also divided *WCS* data into four groups: males starting at 25°C, females starting at 25°C, males starting at 11°C, and females starting at 11°C (Fig. 2B). When flies started at 25°C, males had much a stronger preference for 25°C than female flies. This difference was not observed when they started at 11°C. Moreover, males that started at 25°C strongly preferred 25°C and males that started at 11°C had no preferences between 25°C and 11°C. For females, flies that started at 25°C had an average PI that was higher than those that started at 11°C. But these two groups were not significantly different. Therefore, in the cool avoidance assay, both starting temperatures and genders affect PIs. In the following analysis, to understand the temperature preference of each fly species, males and females were separated and only flies that started at 25°C were analyzed.

### *D. biarmipes, D. bipectinata, D. mojavensis*, and *D. yakuba* don’t avoid warm temperatures

Finally, we calculated PIs of different fly species. As mentioned, only flies that started at 25°C were used. We tested four *D. melanogaster* genotypes: two wild types, *WCS* and *HCS*; a cool receptor mutant, *Ir93a*^*MI*^; and a warm receptor mutant, *Gr28b*^*MB*^ (Knecht et al., 2016, Budelli et al., 2019, Ni et al., 2013). In the warm avoidance assay, both male and female *WCS* and *HCS* strongly preferred 25°C (Fig. 3A,B). PIs of *Gr28b*^*MB*^ were significantly lower than that of *WCS*, which is consistent with previous reports (Ni et al., 2013, Simões et al., 2021, Budelli et al., 2019) (Fig. 3A,B). *D. biarmipes, D. bipectinata*, and *D. yakuba* had similar PIs with *Gr28b*^*MB*^, suggesting that these species don’t have preferences between 25°C and 31°C (Fig. 3A,B). *D. mojavensis* flies had a negative average PI, indicating they prefer 31°C (Fig. 3A,B).

**Fig 3.**
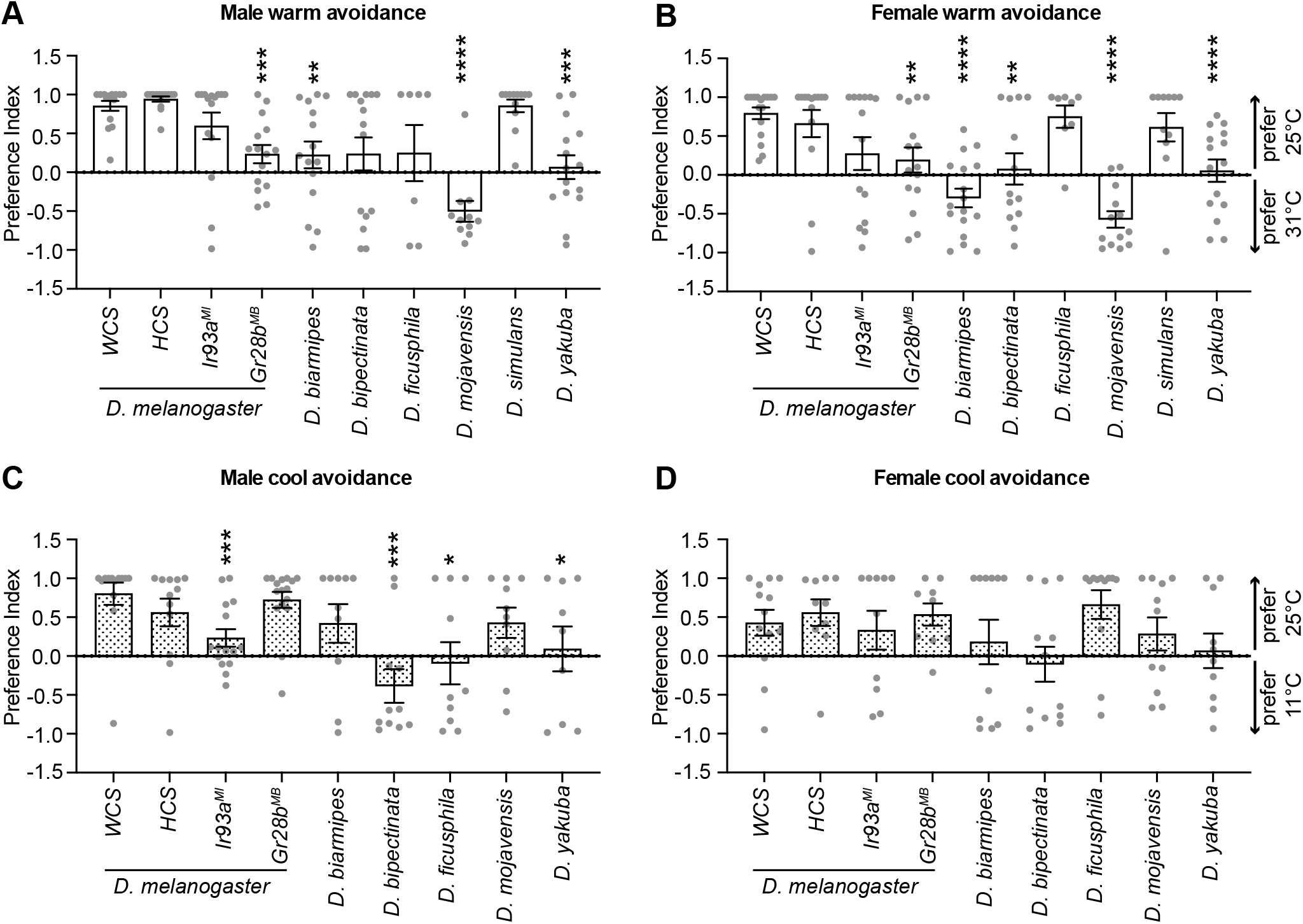
*Drosophila* species have distinct temperature preferences. (A,B) PIs of males (A) and females (B) of indicated *D. melanogaster* genotypes or *Drosophila* species in the warm avoidance assay. (C,D) PIs of males (C) and females (D) of the indicated *D. melanogaster* genotypes or *Drosophila* species in the cool avoidance assay. Data represent mean ± SEM. * *p* < 0.05, ** *p* < 0.01, *** *p* < 0.01 and **** *p* < 0.0001; comparing to the corresponding *WCS*; Mann-Whitney test, except Welch’s test for the comparison of *WCS* and *Gr28b*^*MB*^ and *WCS* and *D. yakuba* in (B).

### *D. bipectinata* and *D. yakuba* don’t avoid cool temperatures

In the cool avoidance assay, male *WCS* and *HCS* strongly preferred 25°C (Fig. 3C). As reported previously (Enjin et al., 2016, Budelli et al., 2019), cool receptor mutant *Ir93a*^*MI*^ males had a lower average PI than *WCS* males (Fig. 3C). Average PIs of *D. ficusphila* and *D. yakuba* males were close to 0, indicating they have no preferences between 25°C and 11°C (Fig. 3C). *D. bipectinata* males had a negative average PI, suggesting they prefer 11°C (Fig. 3C).

Regarding female flies, *WCS* and *HCS* also preferred 25°C (Fig. 3D). Unexpectedly, *Ir93a*^*MI*^ females had a similar average PI with *WCS* females. *D. bipectinata* and *D. yakuba* females, like their males, had PIs close to 0, indicating they have no preferences between 25°C and 11°C (Fig. 3D). On the other hand, *D. ficusphila* females behaved differently from their males: *D. ficusphila* males had no preferences between 25°C and 11°C but their females had strong preferences for 25°C. These data further suggest that genders affect PIs, at least in the cool avoidance assay.

## Discussion

In this study, we modify single-fly thermotactic assays and use TrackMate to track fly movements. We test 801 flies, including four genotypes of *D. melanogaster* and eight other *Drosophila* species. We find that fly species have different temperature preferences from wild-type *D. melanogaster*. Wild-type *D. melanogaster* flies avoid the high temperature of 31°C in the warm avoidance assay and the cool temperature of 11°C in the cool avoidance assay. *D. bipectinata* and *D. yakuba* avoid neither warm nor cool temperatures and *D. biarmipes* and *D. mojavensis* don’t avoid the warm temperature in the warm avoidance assay. Our results also show that starting temperatures and genders affect PIs.

Most fly species move significantly more in warmer environments than in cool environments. But this isn’t true for the cool receptor mutant *Ir93a*^*MI*^ and the warm receptor mutant *Gr28b*^*MB*^ (Knecht et al., 2016, Budelli et al., 2019, Ni et al., 2013). In these two mutants, moving distances are similar in both assays. Moreover, these two mutants move significantly more than wild-type *D. melanogaster WCS* (Fig. 1). Reasons that cause these phenomena are unknown. One possibility is that these mutants *per se* move more. This possibility can be tested by measuring their moving distances in environments with unique temperatures. *Gr28b*^*MB*^ supports this possibility and it moves more than *WCS* in 25°C (Omelchenko et al., 2021). An alternative possibility is that *Ir93a*^*MI*^ and *Gr28b*^*MB*^ move more only when they are allowed to explore different temperature zones. In this case, temperature receptors help animals not only choose an optimal temperature but also save energy. Further studies are needed to test these possibilities.

According to *WCS* data, starting temperatures affect PIs. The only pair that isn’t significantly different is females that start at 25°C and 11°C in the cool avoidance assay. Even in this case, the average PI of the former group is higher than that of the latter group (Fig. 2B). Moreover, genders also affect PIs, at least in the cool avoidance assay. For example, in the cool avoidance assay, *WCS* males have stronger preferences for 25°C than their female counterparts when they start at 25°C (Fig. 2B). In addition, *D. ficusphila* males have no preferences between 25°C and 11°C but females have strong preferences for 25°C (Fig. 3C,D).

GR28BD is the warm receptor that controls warm avoidance when flies are exposed to a steep gradient (Ni et al., 2013, Simões et al., 2021, Budelli et al., 2019). As expected, *Gr28b*^*MB*^ has defects in the warm avoidance assay, but not in the cool avoidance assay (Fig. 3). IR93a is a component of the cool receptor that is required for flies to avoid cool temperatures upon exposure to a steep gradient and its mutant has been reported to be deficient in avoiding both warm and cool temperatures (Enjin et al., 2016, Budelli et al., 2019, Knecht et al., 2016). In our warm avoidance assay, *Ir93a*^*MI*^ flies have lower, but not significantly different PIs compared to *WCS* (Fig. 3A,B). The difference may be because we only analyze flies that start at 25°C. In the cool avoidance assay, *Ir93a*^*MI*^ males have PIs that are significantly lower than *WCS* males (Fig. 3C), which is consistent with previous studies. However, *Ir93a*^*MI*^ females have a similar average PI with *WCS* females (Fig. 3D). We suspect that this is because *WCS* females have lower PIs (Fig. 2B) or our cool avoidance assay uses a lower temperature in the cool zone than the previous study (Budelli et al., 2019). Of note, IR25a is another cool receptor component and its mutant doesn’t have defects in avoiding 10°C (Enjin et al., 2016). Further studies on the functions of the cool receptor are needed.

In the warm avoidance assay, *D. biarmipes, D. bipectinata, D. mojavensis*, and *D. yakuba* show different temperature preferences from *D. melanogaster*. In the cool avoidance assay, *D. bipectinata* and *D. yakuba* have different temperature preferences. Warm and/or cool receptors from these species may offer opportunities to understand how thermoreceptors respond to different temperatures.

In summary, this study uses behavioral assays to understand fly temperature preferences and identifies fly species that have different temperature preferences from *D. melanogaster*. In the future, temperature preferences of other fly species should be analyzed and thermosensory organs, neurons, and molecular receptors should be compared among different fly species to understand mechanisms underlying temperature preferences.

## Materials and methods

### *Drosophila* strains

*White Canton-S* (*WCS*) was used as the wild-type *D. melanogaster* control. *Heisenberg Canton-S* (*HCS*) and *D. mojavensis* were kind gifts from Dr. Michael Dickinson. *Ir93a*^*MI*^ (Knecht et al., 2016) and *Gr28b*^*MB*^ (Ni et al., 2013) were previously reported. Other fly species were from the National *Drosophila* Species Stock Center: *D. ananassae* (14024-0371.11), *D. biarmipes* (14023-0361.03), *D. bipectinata* (14024-0381.21), *D. erecta* (14021-0224.05), *D. ficusphila* (14025-0441.01), *D. simulans* (14021-0251.011), and *D. yakuba* (14021-0261.48).

#### Thermotactic behavioral assay

Flies were raised at 25°C under 12-hour light/12-hour dark cycles and were 3±1 days from eclosure when experiments were performed. All experiments were performed between 8:00 am and 12:00 pm. Fly species that were deemed difficult to distinguish sex via the naked eye were observed and divided under a microscope using a cold plate 24 hours in advance. The warm avoidance assay was performed as described (Omelchenko et al., 2021). Experimental procedures for cool avoidance behavioral assays were identical to previously mentioned procedures, apart from replacing the right-side hot plate with a glass casserole dish filled with ice, with more ice placed on top of the steel plate with the left plate temperature set to 25±1°C and the right plate to 11±1°C.

The images collected from the warm and cool avoidance assay were pre-processed by ImageJ and analyzed by TrackMate as described (Tinevez et al., 2017, Omelchenko et al., 2021). The preference index was calculated by the following formula:

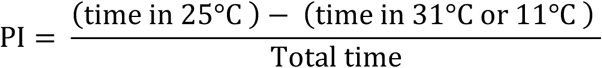

A Python script was developed to calculate moving distances and preference indices.

### Statistical analysis

Statistical details of experiments are mentioned in the figure legends. The normality of distributions was assessed by the Shapiro-Wilk W test (*p* ≤ 0.05 rejected normal distribution). Statistical comparisons of normally distributed data were performed by the Welch’s t test. For data that did not conform to a normal distribution, statistical comparisons were performed by the Mann-Whitney test. Data analysis was performed using GraphPad Prism 9. The pseudo-F statistics was performed by R.

## Acknowledgements

We acknowledge Dr. Jianhong Ou for the R script for the pseudo-F statistics, Dr. Michael Dickinson for *HCS* and *D. mojavensis* flies.

## Data availability

The Python script has been deposited in GitHub and can be accessed at: https://github.com/niflylab/SingleFlyAnalysis.git.

Original statistics and raw data are available at: https://doi.org/10.7910/DVN/DNFWKI.

## Figures

**Movie 1. Set the threshold value for moving distances**. Example trajectories from flies with moving distances of about 250 pixels, 450 pixels, 650 pixels, 850 pixels, and 1050 pixels. Trajectories are shown in red.

